# Biophysical model of axonal stimulation in epiretinal visual prostheses

**DOI:** 10.1101/424622

**Authors:** Michael Beyeler

**Author notes:** M.B. (mbeyeler at uw dot edu) is with the Department of Psychology, the Institute for Neuroengineering, and the eScience Institute at the University of Washington, Seattle, WA 98195.

## Abstract

Visual prostheses aim to restore vision to people blinded from degenerative photoreceptor diseases by electrically stimulating surviving neurons in the retina. However, a major challenge with epiretinal prostheses is that they may accidentally activate passing axon fibers, causing severe perceptual distortions. To investigate the effect of axonal stimulation on the retinal response, we developed a computational model of a small population of morphologically and biophysically detailed retinal ganglion cells, and simulated their response to epiretinal electrical stimulation. We found that activation thresholds of ganglion cell somas and axons varied systematically with both stimulus pulse duration and electrode-retina distance. These findings have important implications for the improvement of stimulus encoding methods for epiretinal prostheses.

## I. INTRODUCTION

Microelectronic retinal prostheses aim to restore vision to people blinded from retinal degenerative diseases by electrically stimulating surviving neurons in the retina. The evoked neuronal responses are transmitted to the brain and interpreted by the patient as visual percepts (‘phosphenes’). Two devices are approved for commercial use: Argus II (epiretinal, Second Sight Medical Products Inc. [1]) and Alpha-IMS (subretinal, Retina Implant AG [2]). In combination with stem cell therapy and optogenetics, various sight restoration options should be available within a decade.

However, a major challenge with current retinal prostheses is the inability to achieve focal tissue activation. Because epiretinal prostheses sit on top of the optic fiber layer, these devices may accidentally stimulate passing axon fibers, which could antidromically activate cell bodies located peripheral to the point of stimulation [3]–[5]. This can cause nontrivial perceptual distortions [6]–[8] that may severely limit the quality of the generated visual experience [9], [10].

Although previous studies have modeled the effect of epiretinal electrical stimulation on the ganglion cell response, most of them either focused on a single neuron (e.g., [3], [11], [12]) or did not consider axons as potential sites of spike initiation (e.g., [11], [13]; but see [14]–[16]). In addition, axonal stimulation has been shown to vary as a function of various stimulus parameters [4], the theoretical foundation of which remains poorly understood.

To examine this issue, we developed a computational model of a small population of morphologically and biophysically detailed retinal ganglion cells, and simulated their response to epiretinal electrical stimulation.

## II. METHODS

### A. Neuron model

The spatial setup of the model is shown in Fig. 1. We populated a ganglion cell layer with a simulated mouse ganglion cell based on 3D morphological data from `neuromorpho.org` (Cell 030102, ID NMO 06380 [17], [18]). The cell’s soma and dendritic tree occupied a 3D box of size 79 µm × 229 µm × 18.7 µm. Starting at the soma, the axon first traveled upward to a distance of *z* ≈ 30 µm, before it bent and traveled along the surface of the simulated optic fiber layer. Ensuring the axon extended well beyond the test area of the extracellular stimulating electrode, we linearly extended the axon by ≈900 µm, starting from the end point of the experimentally traced axon. We then placed a simulated disk electrode (200 µm diameter) at varying distances (100 – 800 µm) above the optic fiber layer.

**Fig. 1:**
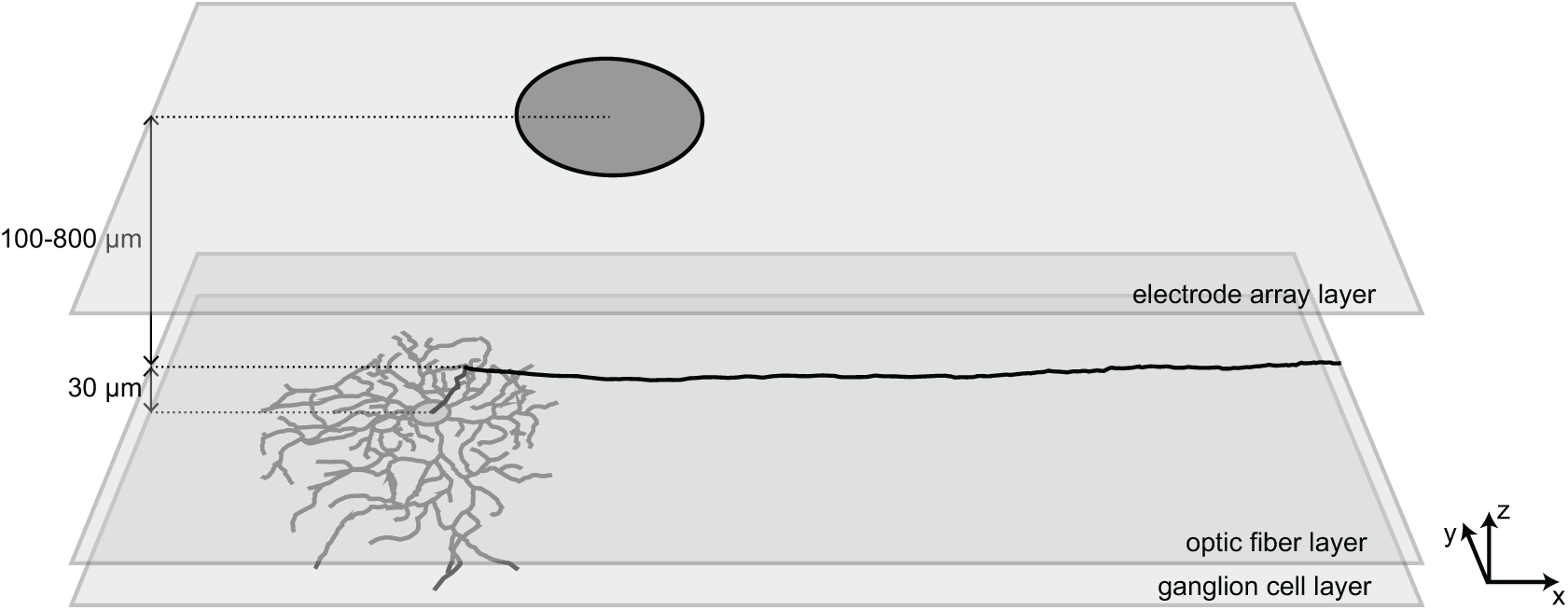
Spatial configuration of the computational model of epiretinal electrical stimulation. A simulated mouse ganglion cell occupied the ganglion cell layer, with its axon extending into the optic fiber layer, and a simulated disk electrode (200 µm diamater) placed at varying distances (100 – 800 µm) above that.

Following [3], [11], [19], we modeled the electrochemical properties of the cell using a Hodgkin-Huxley conductance-based multicompartment model. The soma was modeled as a compartmentalized sphere (24 µm diameter), whereas dendritic tree and axons were split into multiple compartments of varying length (0.1 – 18 µm).

We considered five nonlinear ion channels: Na^+^, Ca^2+^, non-inactivating K^+^ (delayed rectifier), inactivating K^+^ (A type), and Ca^2+^ activated K^+^, as well as a leak current [19]. Following Kirchoff’s law, the membrane potential of each compartment was given as:

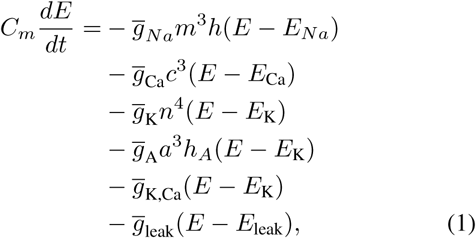

where *C*_*m*_ = 1 µF cm^−2^, *E*_*Na*_ = 35 mV, *E*_Ca_ = 132 mV, *E*_K_ = –75 mV, and *E*_leak_ = − 65 mV. The rate constants for *m, h, c, n, a*, and *h*_*A*_ all solved the first-order kinetic equation:

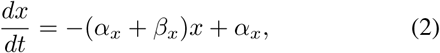

where both *α* and *β* were functions of voltage (see Table I for channel gating kinetics).

**TABLE I:**
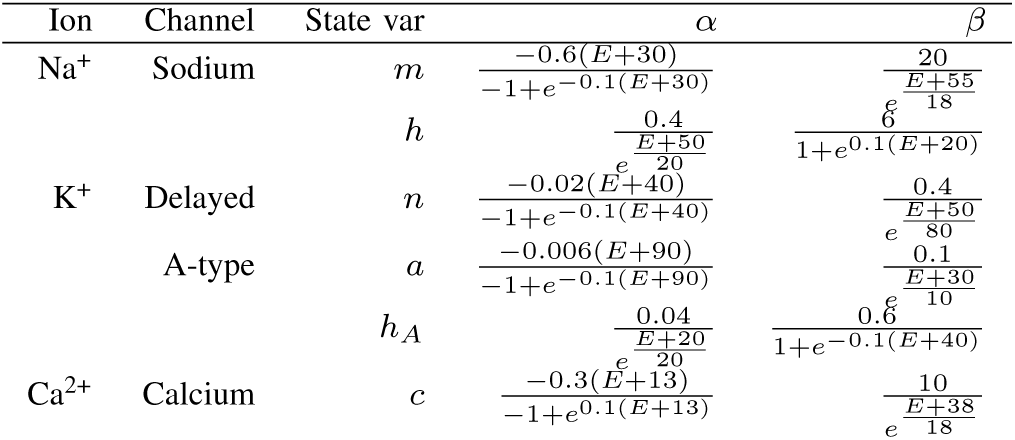
Ion channel kinetics (adapted from [3], [19]).

The gating of *I*_*K,Ca*_ was modeled as:

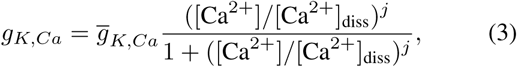

where the Calcium dissociation constant, 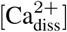, was set to 1 × 10^−6^ M, and *j* was set to 2 [12]. Internal Calcium concentration, [Ca^2+^], was allowed to vary in response to *I*_Ca_. We assumed that the inward flowing Ca^2+^ ions are distributed uniformly throughout the cell and that the free Ca^2+^ above a residual level, [Ca^2+^]_res_ = 1 × 10^−7^ M, was actively removed from the cell with a time constant *τ*_Ca_ = 50 ms:

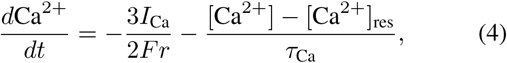

where *F* was the Faraday constant, 3*/r* was the ratio of the surface to volume of the spherical cell soma, and the factor of 2 on *F* was the valency.

Following [3], [19], the five ion channels were distributed with varying densities across dendritic, somatic, and axonal compartments, simulated by varying the value of *g* for each channel (see Table II).

**TABLE II:** Properties of the soma, dendrite, and axon (adapted from [3], [11]).

|  | Dendrites | Soma | Axon hillock | Sodium channel band | Thin segment | Distal axon |
| --- | --- | --- | --- | --- | --- | --- |
| $\bar{g}_{Na}$ (mS cm<sup>-2</sup>) | 25 | 50 – 80 | 100 | 350 – 2800 | 100 – 400 | 70 |
| $\bar{g}_{Ca}$ (mS cm<sup>-2</sup>) | 2 | 1 | 0 | 1.5 | 0 | 0 |
| $\bar{g}_K$ (mS cm<sup>-2</sup>) | 12 | 12 – 18 | 18 | 9 – 72 | 0 – 18 | 18 |
| $\bar{g}_A$ (mS cm<sup>-2</sup>) | 36 | 36 – 54 | 0 | 54 | 0 – 54 | 0 |
| $\bar{g}_{K,Ca}$ (mS cm<sup>-2</sup>) | 0.001 | 0.065 | 0.065 | 0.065 | 0.065 | 0.065 |
| $\bar{g}_{leak}$ (mS cm<sup>-2</sup>) | 0.008 | 0.008 | 0.008 | 0.008 | 0.008 | 0.008 |
| Diameter (μm) |  | 24 | 1 | 1 | 0.4 | 1.0 |
| Length (μm) |  |  | 40 | 40 | 90 | 830 |
| Number of compartments | 2158 | 1 | 19 | 29 | 65 | 705 |

All simulations were run either on an Intel(R) Core(TM) i7-5930K CPU operating at 3.5 GHz using the Brian simulator [20], or an NVIDIA Titan Xp graphics card using the brian2cuda extension. Typically, 10 µs time steps were used. To ensure fine numerical solutions, the ganglion cell was modeled with almost 3000 compartments.

### B. Neural population model

The single cell model was replicated to produce a population of 11 × 6 retinal ganglion cells distributed over a 1000 µm × 500 µm area. To simulate a range of excitation profiles, we uniformly sampled values for 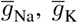, and 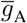 from the naturally occuring range of values reported in [3], [19] (see ranges in Table II).

### C. External stimulus application

External stimulation was given either by an idealized point source or a circular disk electrode.

The electric field *V* (*r*) of a point source was given by:

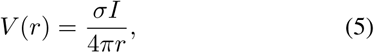

where *σ* = 110 Ω cm was the resistivity of the retinal extracellular solution (typically Ames medium), *I* was the amplitude of the constant current pulse, and *r* was the distance from the stimulating electrode to the point at which the voltage was being computed. Nonuniformities in the electric field arising from the presence of the model cell were not considered.

For stimulation by a circular disk electrode in a semi-infinite homogeneous isotropic half-insulating volume conductor, the extracellular potential was given by [21]:

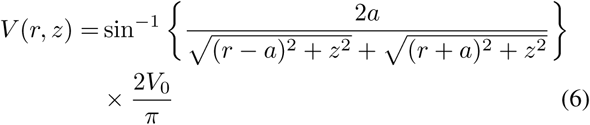

for *z* ≠ 0, where *r* and *z* were the radial and axial distances from the center of the disk, *V*_0_ was the disk potential, *σ* was again the medium conductivity, and *a* = 100 µm. The extracellular potential was calculated at the center of each compartment and uniformly applied to the surface of the entire compartment. The constant voltage model of the disk electrode may be converted to a constant current model (since the extracellular space is modeled as purely resistive) with the addition of a constant multiplicative factor [11], [12].

## III. RESULTS

### A. Excitation thresholds along the axon of a ganglion cell

In physiological experiments with retinal ganglion cells and a small stimulating electrode, the lowest activation thresholds occurred when the electrode was positioned over the sodium channel band (located on the proximal axon, 40 – 80 µm from the soma) [3]. To assess the activation threshold of our model ganglion cell, we applied a 0.2 ms monophasic cathodic pulse originating from a point source (see Eq. 5) located at *z* = 50 µm above the axon. The result is shown in Fig. 2. Similar to rabbit ganglion cells [3], the model cell exhibited its lowest activation threshold when the electrode was centered over the sodium channel band, with thresholds rising across the thin segment and eventually plateauing in the distal axon.

**Fig. 2:**
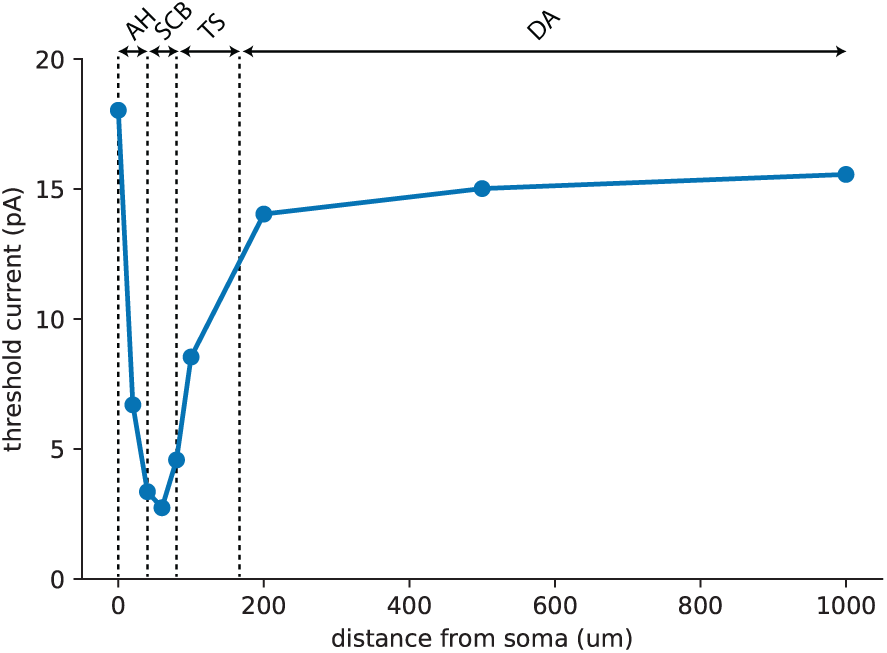
Excitation thresholds along the axon of a multicompartmental Hodgkin-Huxley neuron. AH: axon hillock, SCB: sodium channel band, TS: thin segment, DA: distal axon.

### B. Dynamic range to achieve focal stimulation

The effectiveness with which a stimulus activates only neurons near the electrode can be estimated by studying the relative thresholds for activation of the somatic region (including the axon hillock, sodium channel band, and thin segment) versus activation of the (distal) axon. For example, if thresholds are higher in the distal axon than in the soma, it may be possible to avoid axonal stimulation by utilizing a relatively low stimulus amplitude. We thus measured the range of currents that activated ganglion cell somas without additionally activating axons of passage (‘dynamic range’) by systematically varying the vertical offset (*z*) of an epiretinal disk electrode from the population of simulated ganglion cells. We gradually increased the current amplitude of a 0.2 ms monophasic cathodic pulse until the simulated cells started spiking, and identified the site of spike initiation (i.e., soma versus axon). The resulting excitation thresholds are shown in Fig. 3. Electrode-retina distances below 200 µm tended to activate somas and axons with similar thresholds, thus suggesting that axonal stimulation cannot be avoided at these distances. However, as the vertical offset of the electrode increased, both thresholds and dynamic range increased. The best dynamic range was achieved at an electrode-retina distance of 800 µm.

**Fig. 3:**
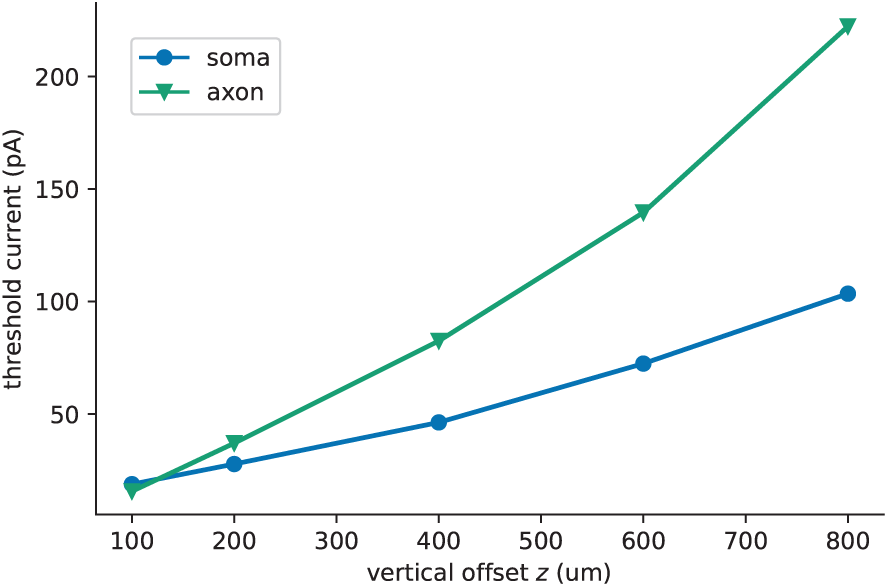
Excitation thresholds of ganglion cell somas and axons in the population model for an epiretinal disk electrode (200 µm diameter) located at varying vertical offsets (*z*).

### C. Response of population model to epiretinal stimulation

A recent electrophysiological study found that stimulus pulse durations two orders of magnitude longer than those typically used in existing epiretinal prostheses can avoid activation of ganglion cell axons [4]. The authors of the study showed that this was due to pulse durations on the order of 10 – 100 ms activating bipolar cells, which in turn activated ganglion cells, thus confining retinal responses to the site of the electrode.

Interestingly, we found a similar effect when we simulated the population response to epiretinal stimulation of 0.1 – 50 ms pulse duration (Fig. 4). Here, we first determined the threshold current for ganglion cells located directly below the electrode. Each dot in Fig. 4 then indicated the number of spikes emitted by a ganglion cell in response to stimulation at 2× threshold. As pulse duration increased, the spatial extent of ganglion cell activation was noticeably reduced.

**Fig. 4:**
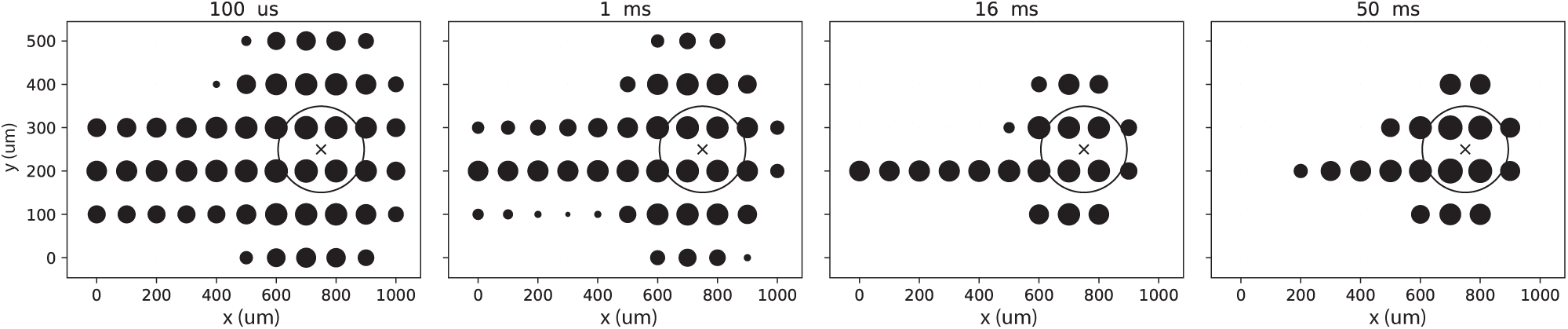
Effect of stimulus pulse duration on ganglion cell responses. Large open circle: epiretinal disk electrode. Small dots: number of spikes emitted by a cell at a particular location (largest dot: 21 spikes). Optic disc lies to the right of each image.

## IV. DISCUSSION

The contributions of this paper are as follows. We found that activation thresholds of ganglion cell somas and axons varied systematically with both stimulus pulse duration and electrode-retina distance. A key prediction of the model is that focal stimulation (i.e., to activate ganglion cell somas without also activating their axons) might be achieved at relatively large electrode-retina distances. However, such distances are also known to give rise to higher activation thresholds and larger phosphene sizes [22]. In addition, the use of longer pulse durations led to a reduction in the spatial extent of axonal activation. A previous study [4] attributed this effect to increased activation of bipolar cells, which in turn activated ganglion cell somas. However, since the present model did not include bipolar cells, the observed effect can be attributed to the spatiotemporal integration properties of the ganglion cell model.

These results suggest that focal activation of the retina might be achieved by a clever combination of stimulus parameters and electrode configuration. In the future, we will extend the model to feature diverse classes of ganglion and bipolar cells. Furthering our understanding of the retinal response to various stimulation patterns and electrode configurations will remain crucial to the continued improvement in the design of epiretinal visual prostheses.

## REFERENCES

1. S. Rizzo, C. Belting, L. Cinelli, L. Allegrini, F. Genovesi-Ebert, F. Barca, and E. diBartolo, “The Argus II retinal prosthesis: 12month outcomes from a single-study center,” American Journal of Ophthalmology, vol. 157, pp. 1282–1290, 2014.

2. K. Stingl, K. U. Bartz-Schmidt, D. Besch, C. K. Chee, C. L. Cottriall, F. Geleker, M. Groppe, T. L. Jackson, R. E. MacLaren, A. Koitschev, and A. K. et al., “Subretinal visual implant Alpha IMS–clinical trial interim report,” Vision Research, vol. 111, pp. 149–160, 2015.

3. J. Jeng, S. Tang, A. Molnar, N. J. Desai, and S. I. Fried, “The sodium channel band shapes the response to electric stimulation in retinal ganglion cells,” Journal of Neural Engineering, vol. 8, pp. 1–12, 2011.

4. A. C. Weitz, D. Nanduri, M. R. Behrend, A. Gonzalez-Calle, R. J. Greenberg, M. S. Humayun, R. H. Chow, and J. D. Weiland, “Improving the spatial resolution of epiretinal implants by increasing stimulus pulse duration,” Science Translational Medicine, vol. 7, pp. 1–11, 2015.

5. T. Esler, R. Kerr, B. Tahayori, D. B. Grayden, H. Meffin, and A. N. Burkitt, “Minimizing activation of overlying axons with epiretinal stimulation: The role of fiber orientation and electrode configuration,” PLOS ONE, vol. 3, 2018.

6. I. Fine and G. M. Boynton, “Pulse trains to percepts: the challenge of creating a perceptually intelligible world with sight recovery technologies,” Philosophical Transactions of the Royal Society B: Biological Sciences, vol. 370, no. 1677, 2015.

7. M. Beyeler, A. Rokem, G. M. Boynton, and I. Fine, “Modeling the perceptual experience of retinal prosthesis patients,” Journal of Vision, vol. 17, pp. 573–573, 2017.

8. M. Beyeler, G. M. Boynton, I. Fine, and A. Rokem, “pulse2percept: A Python-based simulation framework for bionic vision,” in Proceedings of the 16th Python in Science Conference, K. Huff, D. Lippa, D. Niederhut, and M. Pacer, Eds., 2017, pp. 81–88.

9. M. Beyeler, A. Rokem, G. M. Boynton, and I. Fine, “Learning to see again: biological constraints on cortical plasticity and the implications for sight restoration technologies,” Journal of Neural Engineering, vol. 14, no. 5, pp. 1–19, 2017.

10. M. Beyeler, D. Nanduri, J. D. Weiland, A. Rokem, G. M. Boynton, and I. Fine, “A model of ganglion axon pathways accounts for percepts elicited by retinal implants,” bioRxiv, vol. 453035, 2018.

11. R. J. Greenberg, T. J. Velte, M. S. Humayun, G. N. Scarlatis, and J. E. de Juan, “A computational model of electrical stimulation of the retinal ganglion cell,” IEEE Transactions on Biomedical Engineering, vol. 46, no. 5, pp. 505–514, 1999.

12. J. F. Fohlmeister, P. A. Coleman, and R. F. Miller, “Modeling the repetitive firing of retinal ganglion cells,” Brain Research, vol. 510, pp. 343–345, 1990.

13. N. P. Cottaris and S. D. Elfar, “How the retinal network reacts to epiretinal stimulation to form the prosthetic visual input to cortex,” Journal of Neural Engineering, vol. 2, pp. S74–S90, 2005.

14. M. A. Schiefer and W. M. Grill, “Sites of neuronal excitation by epiretinal electrical stimulation,” IEEE Transactions on Neural Systems and Rehabilitation Engineering, vol. 14, no. 1, pp. 5–13, 2006.

15. T. B. Esler, A. N. Burkitt, D. B. Grayden, R. Kerr, B. Tahayori, and H. Meffin, “A computational model of orientation-dependent activation of retinal ganglion cells,” in 38th Annual International Conference of the IEEE Engineering in Medicine and Biology Society (EMBC), 2016, pp. 5447–5450.

16. T. Esler, M. Maturana, R. Kerr, D. B. Grayden, A. N. Burkitt, and H. Meffin, “Biophysical basis of the linear electrical receptive fields of retinal ganglion cells,” Journal of Neural Engineering, vol. 15, 2018.

17. J.-H. Kong, D. R. Fish, R. L. Rockhill, and R. H. Masland, “Diversity of ganglion cells in the mouse retina: Unsupervised morphological classification and its limits,” Journal of Comparative Neurology, vol. 489, no. 3, pp. 293–310, 2005.

18. G. A. Ascoli, D. E. Donohue, and M. Halavi, “NeuroMorpho.Org: a central resource for neuronal morphologies,” Journal of Neuroscience, vol. 27, no. 35, pp. 9247–9251, 2007.

19. B. W. Sheasby and J. F. Fohlmeister, “Impulse encoding across the dendritic morphologies of retinal ganglion cells,” Journal of Neurophysiology, vol. 81, pp. 1685–1698, 1999.

20. D. Goodman and R. Brette, “The Brian simulator,” Frontiers in Neuroscience, vol. 3, p. 26, 2009.

21. J. D. Wiley and J. G. Webster, “Analysis and control of the current distribution under circular dispersive,” IEEE Transactions on Biomedical Engineering, vol. BME-29, pp. 381–385, 1982.

22. D. Nanduri, I. Fine, A. Horsager, G. M. Boynton, M. S. Humayun, R. J. Greenberg, and J. D. Weiland, “Frequency and amplitude modulatoin have different effects on the percepts elicited by retinal stimulation,” Investigative Ophthalmology and Visual Science, vol. 53, no. 1, pp. 205–214, 2012.

